# Brain Preparedness for Active Forgetting: Cortisol Awakening Response Proacts Prefrontal Control Over Hippocampal-Striatal Circuitry

**DOI:** 10.1101/2025.09.14.673810

**Authors:** Bingsen Xiong, Changming Chen, Yucheng Bian, Yanqiu Tian, Haowen Su, Manqi Sha, Shouwen Zhang, Chao Liu, Jianhui Wu, Shaozheng Qin

**Affiliations:** Beijing Key Laboratory of Applied Experimental Psychology, National Demonstration Center for Experimental Psychology Education, Faculty of Psychology, Beijing Normal University, Beijing 100875, China; School of Education, Chongqing Normal University, Chongqing, 401331, China; State Key Laboratory of Cognitive Neuroscience and Learning & IDG/McGovern Institute for Brain Research, Beijing Normal University, Beijing 100875, China; West Essence Clinic, Beijing Institute of Functional Neurosurgery & Xuanwu Hospital, Capital Medical University, Beijing 100053, China; Shenzhen Key Laboratory of Affective and Social Cognitive Science, Shenzhen University, Shenzhen 518060, China; Chinese Institute for Brain Research, Beijing 100069, China

**Keywords:** memory suppression, cortisol awakening response, dorsolateral prefrontal cortex, hippocampus, caudate

## Abstract

Stress-related psychiatric pathologies (e.g., rumination in depression, intrusive memories in PTSD) involve impaired inhibitory control over emotional memories and disrupted cortisol circadian rhythms. However, how these mechanisms interact to support adaptive forgetting remains unclear. Here, by integrating multi-modal neuroendocrine profiling with fMRI-based memory suppression in 90 participants, we demonstrate that the cortisol awakening response (CAR)—a circadian surge priming the brain for daily demands—proactively and selectively enhances suppression of recently acquired but not overnight-consolidated emotional memories. This time-dependent facilitation manifests through dual neural mechanisms: Robust CAR broadly amplifies dorsolateral prefrontal cortex (dlPFC) engagement during suppression attempts and specifically strengthens top-down dlPFC control over hippocampal-striatal circuits exclusively for recent traces. Our findings bridge circadian biology, systems neuroscience, and clinical psychopathology to establish CAR as a neuroendocrine preparedness mechanism that proactively configures adaptive forgetting circuits across sleep-wake cycles, thereby informing circadian-based interventions for stress disorders.

## 1. Introduction

The ability to actively forget unwanted emotional memories is crucial for maintaining mental health and ensuring survival (Anderson & Hulbert, 2021). Dysfunctions in this memory control system have been linked to symptoms in various psychiatric disorders, such as rumination in depression and intrusive memories in posttraumatic stress disorder (PTSD) (Liu et al., 2016). Notably, stress-related mental disorders characterized by impaired active forgetting often concurrently exhibit atypical circadian rhythms of the stress hormone cortisol (Kalafatakis et al., 2018; Kondratova & Kondratov, 2012; Menet & Rosbash, 2011; Nassan & Videnovic, 2022; Oster et al., 2017), suggesting a conceivable assumption that cortisol’s circadian activity may play a critical role in modulating active forgetting of aversive memories. Indeed, the circadian system is well-recognized to generally influence numerous psychophysiological processes, including sleep, alertness and cognitive functions (Nassan & Videnovic, 2022); cortisol activity serves as a key modulator of memory systems through nongenomic and relatively slower genomic actions on hippocampal and prefrontal-limbic networks (de Kloet, de Kloet, de Kloet, & de Kloet, 2019). However, the specific mechanisms by which cortisol circadian activity modulate active forgetting of unwanted emotional memories, and the underlying neural substrates remain largely unexplored.

The cortisol awakening response (CAR), a crucial feature of cortisol circadian rhythm, is a candidate for modulating adaptive control of emotional memories proactively. Characterized by a significant rise in cortisol levels within 30 to 40 minutes after waking from nighttime sleep, the CAR is thought to prepare the brain and cognitive systems to maintain homeostasis and promote adaptive responses for the upcoming day (Clow, Thorn, Evans, & Hucklebridge, 2004; Elder, Wetherell, Barclay, & Ellis, 2014; Fries, Dettenborn, & Kirschbaum, 2009; Law, Hucklebridge, Thorn, Evans, & Clow, 2013; Stalder et al., 2025). Beyond evidence supporting this preparation hypothesis from behavioral studies (Adam, Hawkley, Kudielka, & Cacioppo, 2006; Kunz-Ebrecht, Kirschbaum, Marmot, & Steptoe, 2004; Law et al., 2020; Schlotz, Hellhammer, Schulz, & Stone, 2004; Stalder, Evans, Hucklebridge, & Clow, 2010a, 2010b), recent pharmacological fMRI studies further established a causal link between the CAR and its proactive role in priming optimal prefrontal-hippocampal functional organization involved in executive control and emotional processing essential for active forgetting of aversive memory (Chen et al., 2025; Xiong et al., 2021; Zeng et al., 2024). However, direct evidence is lacking to support the CAR-mediated regulation of active forgetting emotional memories. Critically, how CAR specifically modulates the suppression efficacy of memories at different consolidation stages remains unexplored.

Recent advances in human cognitive neuroscience have expanded our understanding of the neurocognitive mechanisms underlying active forgetting, underscoring a top-down inhibitory control that is exerted by the prefrontal cortex over mnemonic activity in the hippocampus among other regions. This process is characterized by prominent activation in the dorsolateral prefrontal cortex (dlPFC) and concurrent disengagement of the hippocampus (Anderson & Hanslmayr, 2014; Anderson & Hulbert, 2021; Catarino, Kupper, Werner-Seidler, Dalgleish, & Anderson, 2015; Walker & Stickgold, 2004; Walker & van der Helm, 2009). Moreover, memories become more resistant to suppression following overnight consolidation, necessitating stronger prefrontal engagement and more pronounced disengagement of the hippocampus (Liu et al., 2016). Emerging research further highlights that the striatum— particularly the caudate—acts as a supramodal hub for inhibitory control. It interacts with the dlPFC through prefrontal-striatal projections to suppress hippocampal retrieval via indirect pathway activation (Anderson & Hulbert, 2021; Burke, Rotstein, & Alvarez, 2017; Depue, Orr, Smolker, Naaz, & Banich, 2016; Guo, Schmitz, Mur, Ferreira, & Anderson, 2018; R. Levy, Friedman, Davachi, & Goldman-Rakic, 1997; Robbins & Cools, 2014). Integrating these mechanisms with the preparatory effects of the CAR, we hypothesize that the CAR-mediated proactive tone would synergize with voluntary suppression efforts, thereby enhancing prefrontal-originated top-down inhibitory control over the hippocampal-striatal circuits, thus promoting active forgetting of emotional memories.

To test this hypothesis, we designed a multimodal protocol integrating synchronized neuroendocrine profiling with memory suppression and overnight consolidation paradigms (**Fig. 1A**). This experimental design, in conjunction with a tripartite analytical framework that integrates univariate activation, functional connectivity and dynamic causal modeling (DCM), allows us to test three dissociable sub-hypotheses: (**H1**) Behaviorally, we hypothesized that a stronger CAR would selectively enhance suppression-induced forgetting for recently encoded emotional memories, but not for memories that had undergone overnight consolidation— suggesting a time-sensitive modulation of suppression efficacy by CAR. (**H2**) At the regional activation level, we expected that higher CAR would predict increased recruitment of the dlPFC alongside reduced activation in the posterior hippocampus and striatum, particularly during suppression of recent memories—reflecting CAR’s priming role in modulating neural substrates of inhibition. (**H3**) At the network level, we hypothesized that greater CAR would be associated with stronger top-down control from the dlPFC to hippocampal and striatal targets, as indexed by enhanced functional and effective connectivity specifically during suppression of recent memories. Together, these hypotheses integrate circadian biology, cognitive neuroscience, and clinical frameworks to propose a chronotherapeutic model in which phase-specific enhancement of the CAR may modulate fronto-hippocampal-striatal dynamics to reduce vulnerability to intrusive memories and maladaptive emotions—a potential mechanism for refining intervention strategies for stress-related disorders.

**Fig. 1.**
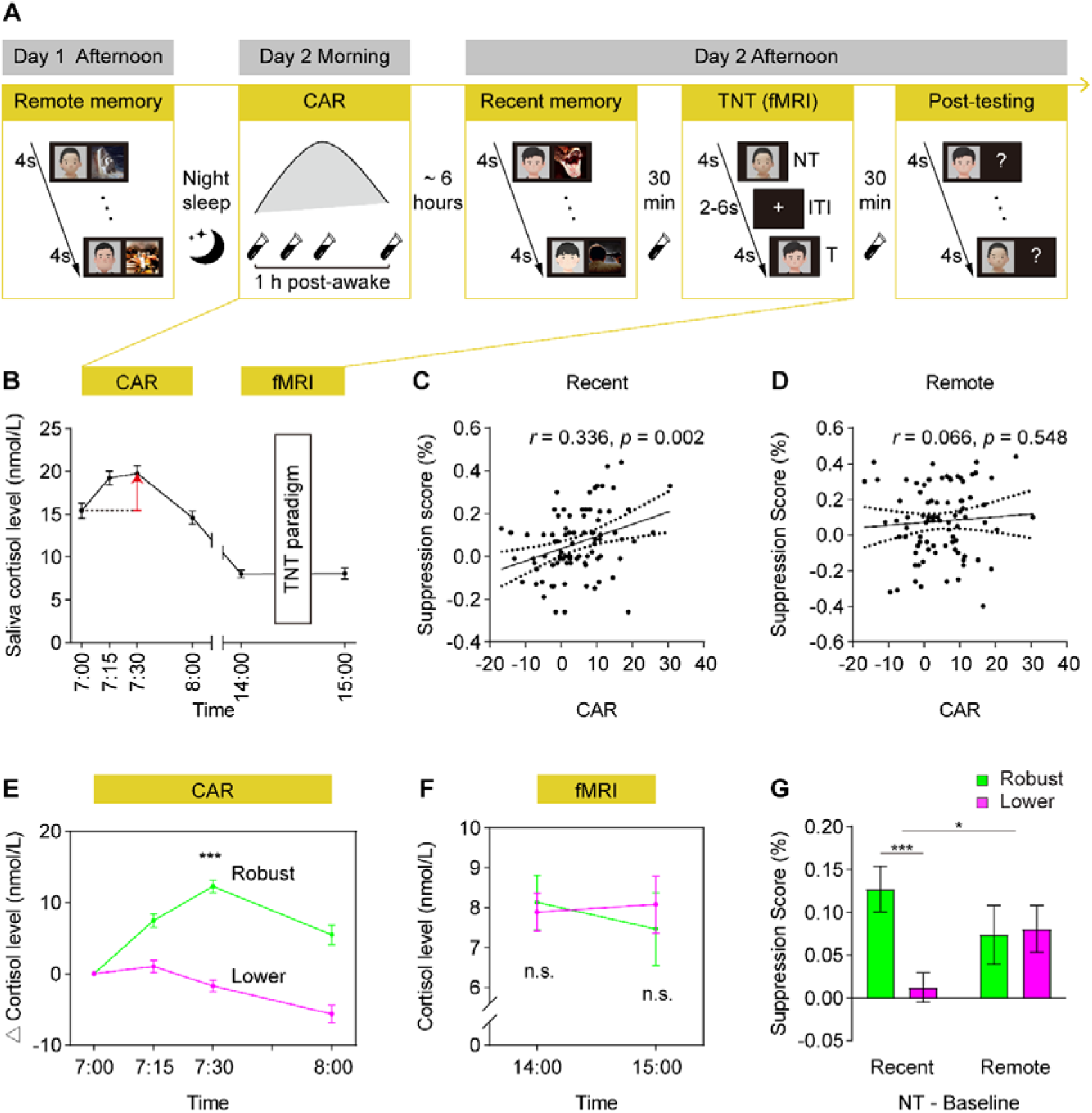
Experimental design and proactive effects of the CAR on active suppression of emotional memory. (**A**) The experiment consisted of five phases. Participants performed two memory acquisition sessions on Day 1 (Remote memory) and Day 2 (Recent memory) afternoon, which occurred 24 hours and 30 minutes before the Think/No-Think (TNT) phase, respectively. Participants were trained to memorize 26 face-scene pairs in each of the two acquisition sessions. During the TNT phase, participants underwent fMRI scanning while they were performing a ‘Think/No-Think’ task. During the Post-testing phase, participants were given faces as cues and asked to recall their associated target pictures. Overall **CAR** profile and diurnal cortisol levels were assessed at 4-time points within 1-hour immediately post-wakening as well as 2-time points right before and after fMRI scanning on Day 2. (**B**) Salivary cortisol levels at 4-time points (i.e., 0-, 15-, 30- and 60-minute) within 1 hour after awakening in the morning, as well as at 2-time points right before and after fMRI scanning in the afternoon when participants performed a TNT task. (**C-D**) Scatter plots illustrate positive correlation of individual’s CAR with Recent but not Remote memory suppression score. (**E-F**) Cortisol levels of individuals with Robust- and Lower-CAR in the morning, before and after fMRI scanning in the afternoon. (**G**) Bar graphs depict higher suppression scores in individuals with Robust-than Lower-CAR only in Recent but not Remote condition about 6 hours later in the afternoon of the same day. Notes: To protect personal privacy and avoid disclosure of identifying information, we have replaced the original real experimental face-related materials with computer-generated face images; n.s., not significant; *, *p* < 0.05; **, *p* < 0.01; ***, *p* < 0.001; Error bars represent standard error of mean.

## 2. Materials and Methods

### 2.1 Participants

A total of 90 young, healthy, male college students participated in this study (mean age: 22.06 ± 1.33 years; age range: 18-26 years). Only male participants were included to avoid confounding effects from hormonal fluctuations during the menstrual cycle and the potential influence of hormonal contraceptives in young adult females (Cousijn et al., 2010; Kirschbaum, Kudielka, Gaab, Schommer, & Hellhammer, 1999). All participants reported no history of neurological, psychiatric, or endocrine disorders. Exclusion criteria comprised current medication that affects the neural or endocrine systems, daily tobacco or alcohol use, irregular sleep-wake patterns, intense daily physical exercise, abnormal hearing or uncorrected vision, predominant left-handedness, active periodontitis (Mathew et al., 2019), and recent stressful experiences or major life events (see *Supplementary Materials* for more details).

Data from 4 participants were excluded from the analyses due to incomplete salivary samples or suboptimal behavioral performance (beyond ±3 standard deviation). Consequently, a final sample of 86 participants was included in the behavioral data analysis. An additional 15 participants were excluded due to excessive head movement during brain scanning (exceeding 3 mm/degree), resulting in 67 participants being included in the subsequent brain imaging analyses (*Table S1*).

After excluding 4 participants due to incomplete salivary samples or behavioral outliers (> ±3SD), 86 entered behavioral analysis. Subsequently, 15 were excluded for excessive head motion (>3mm/degree), yielding 67 for neuroimaging analyses (*Table S1*).

Informed written consent was obtained from all participants prior to the experiment, and the study protocol was approved by the Institutional Review Board for Human Subjects at Beijing Normal University.

### 2.2 General Experimental Procedure

The experimental protocol comprised five phases (**Fig. 1A**). Memory acquisition was divided into two sessions: the first on Day 1 and the second on Day 2, which occurred 24 hours (Remote condition) and 30 minutes (Recent condition) before fMRI scanning, respectively. On Day 1, participants underwent training to learn and retain a set of 26 face-scene picture associations. On Day 2, they received additional training to memorize another set of 26 face-scene associations. 30 minutes after the second training session, participants engaged in a memory suppression task using the modified ’Think/No-Think (TNT)’ paradigm during fMRI scanning. Subsequently, participants completed a post-scan memory test for face-scene associations to evaluate their memory performance and the effectiveness of memory suppression. Salivary samples were collected at six-time points on Day 2 to assess the CAR and diurnal cortisol rhythms.

### 2.3 Physiological and Psychological Measures

#### Salivary cortisol measurement

Cortisol levels were quantified from six saliva samples per participant using Salivette tubes (Sarstedt, Germany). Samples were collected at four timepoints post-awakening (0, 15, 30, and 60 minutes) and during fMRI scanning (pre- and post-session) on Day 2 (**Fig. 1**). Participants abstained from eating or drinking 1 hour prior to sampling and were restricted from consuming alcohol, caffeine, and nicotine, as well as engaging in strenuous exercise for 24 hours before the procedure. To ensure participant adherence and maintain data quality, four specific strategies were implemented (Stalder et al., 2016; Tian et al., 2021; Wu et al., 2015; Zhu et al., 2019): 1) Individual pre-experiment instruction; 2) Pre-scheduled wake-up and sampling alarms; 3) Time-stamped photographs of sample collections; 4) Electronic monitoring of sampling times (for further details, see *Supplementary Materials*). Samples were stored at -20°C until analysis, then centrifuged at 3000 rpm for 5 minutes, and assayed using an electrochemiluminescence immunoassay (ECLIA; Cobas e601, Roche Diagnostics, Mannheim, Germany). The assay demonstrated a detection range of 0.5-1750 nmol/L with intra- and inter-assay coefficients of variation < 10%. The CAR was calculated as the increment from 0 to 30 minutes post-awakening (R30) (Clow et al., 2004; Stalder et al., 2016).

#### Stimuli

The study utilized 52 face-scene pairs consisting of standardized stimuli selected through rigorous protocols. Face stimuli (26 male/26 female) were drawn from a pool of 100 Chinese portrait photographs based on predefined criteria: neutral expression with direct gaze (valence = 5.16 ± 0.53, arousal = 5.03 ± 0.43; validated using a 9-point scale in pilot testing), absence of accessories (e.g., headdress, glasses, beards), and equivalent ratings across genders for arousal, valence, attractiveness, and trustworthiness (all *p* > 0.05). Scene stimuli (52 items) were sourced from the International Affective Picture System and characterized by high negative valence (2.37 ± 0.69) and arousal (7.89 ± 0.55), with minimal semantic overlap ensured. Face and scene images were randomly paired across participants to create 52 face-scene associations, which were subsequently randomized for assignment to Day 1 and Day 2.

#### Memory acquisition

During the memory acquisition phase (**Fig. 1A**), participants underwent training sessions on Day 1 and Day 2, conducted outside the scanner. These sessions occurred 24 hours and 30 minutes before the TNT task, respectively. To minimize variability, the time interval between sessions was strictly controlled. Training began at 14:00 on Day 1, with scanning scheduled for 14:00–15:00 on Day 2, and Day 2 training commenced 30 minutes prior to scanning. In each session, participants memorized 26 face-scene picture pairs through multiple study-recall cycles. During these cycles, each association was displayed for 4 seconds, and participants were instructed to remember the details. After all associations were presented, participants viewed a face and recalled the corresponding scene. This process was repeated 2–4 times until all 26 associations were accurately recalled. Participants provided detailed descriptions of the associated images when cued by faces, ensuring vivid episodic memories rather than vague impressions. Individuals requiring more than 4 cycles to achieve 100% accuracy were excluded to prevent overly strong memories that might resist suppression.

#### Memory Suppression

During the memory suppression phase (**Fig. 1A**), participants underwent fMRI scanning while performing the TNT task for face-scene associations acquired either 24 hours prior (Remote) or 30 minutes prior (Recent). Each trial began with the presentation of a face stimulus for 4 seconds, followed by an inter-trial interval (ITI) during which a fixation cross was displayed for 2–6 seconds (average duration: 4 seconds). The ‘Think’ and ‘No-Think’ trials were pseudo-randomized across participants and interleaved with fixation periods. Instructional cues—green rectangles for ‘Think’ trials and red rectangles for ‘No-Think’ trials—appeared simultaneously with the face stimuli, guiding participants to either recall the associated scene (‘Think’) or actively suppress it from entering consciousness (‘No-Think’). In this phase, only faces were presented, ensuring that participants manipulated memories associated with the target scenes. After the presentation of each face, a fixation cross appeared during the ITI, serving as a baseline for experimental trials. The entire task lasted 19.2 minutes and consisted of 144 trials in total, which were evenly distributed across four conditions (36 trials per condition). Participants viewed 36 out of 52 unique faces, half of which were learned on Day 1 (Remote condition) and the other half on Day 2 (Recent condition). An additional 16 trials served as the baseline condition and were not included in this phase. Half of the faces were randomly assigned to either the ‘Think’ or ‘No-Think’ condition, resulting in a 2-by-2 full factorial design with four experimental conditions: memory suppression (No-Think vs. Think) crossed with acquisition time (Remote vs. Recent). Each face was repeated four times, yielding a total of 144 trials. Participants were explicitly instructed to directly suppress unwanted memories during ‘No-Think’ trials.

#### Memory testing

During the testing phase (**Fig. 1A**), memory performance for face-scene associations was evaluated using a cued-recall task that included all 52 faces learned on Day 1 and Day 2. Each face served as a cue, prompting participants to verbally recall and describe the associated scene in as much detail as possible within a maximum of 30 seconds per trial. Responses were scored as correct only if they contained sufficient unique details to clearly identify the specific scene. Descriptions lacking clarity or accuracy were classified as forgotten. Three independent raters, who were blinded to the experimental conditions, reviewed the responses. Final test scores were determined through cross-validation among the three raters. A consensus was required for each item’s final judgment; any disagreements were resolved through discussion among the raters.

#### Questionnaires

Upon arrival at the laboratory, participants were administered two questionnaires: the State-Trait Anxiety Inventory (STAI) (Spielberger, 1985), which assesses both state and trait anxiety levels, and the Perceived Stress Scale (PSS) (Cohen, 1988), which evaluates long-term psychological stress. Additionally, a sleep questionnaire was utilized to monitor participants’ sleep quality throughout the experimental period (see Supplementary Materials for details).

### 2.4 Behavioural Data Analysis

To verify the effectiveness of manipulation in the TNT task, memory accuracy was analyzed using a 2-by-3 repeated-measures analysis of variance (ANOVA), with memory suppression (Baseline vs. Think vs. NoThink) and acquisition time (Recent vs. Remote) as within-subject factors (*Fig. S1*). Suppression scores for overnight (Remote) and recently acquired (Recent) memories were calculated by subtracting the memory accuracy of NoThink items from their corresponding Baseline items, providing a measure of suppression-induced forgetting or suppression efficiency (B. J. Levy & Anderson, 2012; Liu et al., 2016).

To investigate how individual differences in the CAR modulate suppression-induced forgetting, we performed multiple regression analyses separately for Recent and Remote memory suppression scores. In these analyses, CAR served as the covariate of interest, while sleep duration, perceived stress, and state-trait anxiety were treated as covariates of no interest. To further explore the interaction effect between CAR and suppression scores, we conducted complementary analyses by dividing participants into two groups: those with a Robust-CAR and those with a Lower-CAR. Following the criteria established in prior studies (Clow et al., 2004; Wust et al., 2000; Xiong et al., 2021), individuals whose cortisol levels increased by more than 50% within 30 minutes after awakening were classified into the Robust-CAR group, whereas those with less than a 50% increase were assigned to the Lower- CAR group. We first confirmed the group difference in CAR by conducting an independent sample t-test on R30 (i.e., the rise in cortisol level from awakening to 30 minutes post-awakening). Subsequently, we performed a 2-by-2 repeated-measures ANOVA on the suppression score, with acquisition time (Recent vs. Remote) as the within-subject factor and group (Robust-CAR vs. Lower-CAR) as the between-subject factor.

### 2.5 Brain Imaging Data Acquisition

Whole-brain images were acquired using a Siemens 3.0 Tesla TRIO MRI scanner (Erlangen, Germany) at the National Key Laboratory of Cognitive Neuroscience and Learning & IDG/McGovern Institute for Brain Research, Beijing Normal University. Functional brain images were collected during the TNT task with a gradient-recalled echo planar imaging (GR-EPI) sequence (33 slices, volume repetition time = 2.0 s, echo time = 30 ms, flip angle = 90°, slice thickness = 4 mm, gap = 0.6 mm, field of view = 200 × 200 mm, voxel size = 3.1 × 3.1 × 4.0 mm). High-resolution anatomical images were obtained using a T1-weighted 3D magnetization-prepared rapid gradient echo sequence (192 slices, volume repetition time = 2530 ms, echo time = 3.45 ms, flip angle = 7°, slice thickness = 1 mm, field of view = 256 × 256 mm, voxel size = 1 × 1 × 1 mm³).

### 2.6 Brain Imaging Data Analysis

#### 2.6.1 Preprocessing

Image preprocessing and statistical analysis of fMRI data were conducted using Statistical Parametric Mapping (SPM12, http://www.fil.ion.ucl.ac.uk/spm). The first four functional image volumes were discarded to allow for signal stabilization and participant adaptation to scanning noise. The remaining images underwent slice-timing correction, followed by realignment to correct for head motion artifacts. These images were then co-registered to the gray matter segment derived from anatomical T1-weighted images and spatially normalized into the standard Montreal Neurological Institute (MNI) stereotactic space. Subsequently, the images were resampled into 2-mm isotropic voxels and smoothed with a three-dimensional isotropic Gaussian kernel of 6 mm full-width at half-maximum (FWHM).

#### 2.6.2 Univariate GLM Analysis

The data were statistically analyzed within the framework of general linear models (GLM). Separate regressors of interest were modeled for the four experimental conditions (memory suppression: Think vs. NoThink; acquisition time: Recent vs. Remote) and convolved with the canonical hemodynamic response function (HRF) implemented in SPM12 at the first level. Additionally, the six parameters accounting for head motion were included as covariates to control for movement-related variability. The analysis incorporated a high-pass filter with a cutoff of 1/40 Hz and applied a serial correlation correction using a first-order autoregressive model (AR(1)). Parameter estimate images reflecting relevant contrasts were initially generated at the individual subject level and subsequently entered into a group-level analysis, where participants were treated as a random effect.

To evaluate the neural activity associated with memory retrieval (Think) and suppression (NoThink) for recently acquired (Recent) and overnight consolidated (Remote) memories, we performed a 2-by-2 repeated-measures ANOVA at the whole-brain level with acquisition time (Recent vs. Remote) and TNT (Think vs. NoThink) served as within-subject factors. Significant clusters were identified using a height threshold of *p* < 0.001 and an extent threshold of *p* < 0.05 with cluster-based FWE correction. Subsequently, ANOVAs were conducted on data extracted from these significant clusters, and the Time-by-TNT interaction effects were visualized in bar graphs.

To examine how individual differences in the CAR modulate brain activity associated with memory suppression, we conducted whole-brain multiple regression analyses separately for Recent and Remote memory suppression. In these analyses, the individual CAR was treated as the covariate of interest, while sleep duration, perceived stress, and state-trait anxiety were included as nuisance covariates. Significant clusters were identified using the same criterion described above. Correlation analyses were then performed on data extracted from these clusters, and the patterns of correlation were visualized using scatterplots.

To further investigate the interaction effects of CAR on brain function during memory suppression, we performed a complementary 2-by-2 repeated measures ANOVA at the whole-brain level, with acquisition time (Recent vs. Remote) as the within-subject factor and group (Robust-CAR vs. Lower-CAR) as the between-subject factor. Significant clusters were identified using the same threshold criterion outlined above. Subsequent ANOVAs were performed on data extracted from these significant clusters, and the main effect of Group as well as the Group-by-Time interaction effect were visualized in bar graphs.

#### 2.6.3 Task-Dependent Functional Connectivity Analysis

To examine whether individual differences in the CAR modulate functional coupling among brain systems involved in memory suppression, we conducted separate generalized psychophysiological interaction (gPPI) analyses (McLaren, Ries, Xu, & Johnson, 2012), using clusters identified in the above univariate analyses as seeds of interest. The mean time series extracted from these regions of interest (ROIs) were deconvolved to estimate underlying neuronal activity (i.e., the physiological variable). Subsequently, four PPI regressors corresponding to each task regressor at the individual level were generated by multiplying the estimated neuronal activity from the seed region with a vector coding for the effects of each condition, resulting in four psychophysiological interaction vectors. These vectors were further convolved with a HRF to produce four PPI regressors of interest. Additionally, task-related activations were included in the general linear model (GLM) to account for and remove the effects of common driving inputs on brain connectivity.

Contrast images reflecting PPI effects at the individual level were subsequently entered into group-level analyses. Specifically, we performed whole-brain multiple regression analyses separately for suppression of Recent and Remote memory, with CAR as the covariate of interest, while controlling for sleep duration, perceived stress, and state-trait anxiety as nuisance covariates. Significant clusters were identified using the same statistical threshold described above. Correlation analyses were then conducted on data extracted from these clusters, and the resulting correlation patterns were visualized in scatterplots. Additionally, we carried out 2-by-2 ANOVAs at the whole-brain level, with acquisition time (Recent vs. Remote) as a within-subject factor and group (Robust-CAR vs. Low-CAR) as a between- subject factor. Significant clusters were again determined using the aforementioned threshold. Subsequent ANOVAs were performed on data extracted from these clusters, and the Group- by-Time interaction effects were illustrated in bar graphs.

#### 2.6.4 Dynamic Causal Modelling

To further investigate how CAR modulates directed causal influences within prefrontal- hippocampal-striatal circuits during active forgetting of emotional memory, we employed dynamic causal modeling (DCM) supplemented by a hierarchical Parameter Empirical Bayes (PEB) framework (Friston, Harrison, & Penny, 2003; Zeidman, Jafarian, Corbin, et al., 2019; Zeidman, Jafarian, Seghier, et al., 2019). Within DCM architecture, neural state dynamics are mathematically described through differential equations that characterize the temporal evolution of regional neural activity (ż) as a function of experimental inputs (u) and inter/intra-regional connectivity:

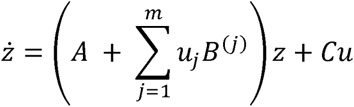

The experimental inputs enter the model in two ways: driving inputs that directly activate specific nodes at stimulus onset, and modulatory inputs that alter connection strengths between nodes. These dynamics are formalized through three connectivity matrices: (1) the A-matrix specifies the intrinsic connectivity within the network of brain regions; (2) the B- matrix captures changes in effective connectivity induced by modulatory inputs; (3) and the C-matrix quantifies the rate of change in the neural response due to driving inputs. Furthermore, each brain region is endowed with an inhibitory self-connection, which reflects its gain or sensitivity to incoming inputs.

Individual-level analysis commenced with ROI selection based on aforementioned gPPI results, identifying four key regions: bilateral dlPFC, left hippocampus, and right caudate. For each participant, we extracted the principal eigenvariate time series from these ROIs under NoThink_Recent (i.e., suppressing recently acquired memories) condition. A full DCM architecture was then specified (**Fig. 4A**), incorporating bidirectional connections between all ROIs in the A-matrix; inhibitory self-connections for gain regulation; NoThink_Recent as driving input (C-matrix) to all nodes; NoThink_Recent-induced modulatory effects (B-matrix) on all connections.

Group-level analysis employed PEB framework to integrate individual DCM parameters while accounting for inter-subject variability (Zeidman, Jafarian, Seghier, et al., 2019). This hierarchical Bayesian approach estimates inter-group effects through maximizing model evidence (quantified by free energy F), achieving an optimal balance between model fitness and complexity. Higher *F* values indicate stronger evidence for models that appropriately reconcile explanatory power with parsimony. In the present study, we aimed to identify the most plausible explanation for the group differences (Robust-CAR vs. Lower-CAR) in terms of changes in effective connectivity, modulated by memory suppression. Starting with the full model, we systematically defined a model space comprising reduced models with varying configurations of the B-matrix. Specifically, in each reduced model, the modulatory effect of memory suppression on the connection between two regions was switched off. Overall, the model space consisted of 12 candidate models: one full model, ten reduced models, and one null model without any modulation, serving as a baseline (**Fig. 4B**). Bayesian Model Comparison (BMC) was conducted to evaluate relative model evidence, followed by Bayesian Model Averaging (BMA) to derive robust parameter estimates across models (Penny, Flandin, & Trujillo-Barreto, 2007).

## 3. Results

### 3.1 CAR Selectively Enhances Suppression-Induced Forgetting of Recent Emotional Memories

We first evaluated the characteristics of the CAR profile and diurnal cortisol levels. As shown in **Fig. 1B**, participants’ cortisol response peaked at around 30 minutes post-awakening, followed by a gradual decline at 60 minutes, and maintained a relatively low and stable level spanning over the afternoon (*F*_5, 510_ = 45.931, *p* < 0.001). The overall CAR was quantified by an increase in cortisol levels within 30 minutes post-awakening relative to immediately after awakening, namely R30 in literature(Stalder et al., 2016).

To examine whether the CAR modulates the effectiveness of suppressing recently acquired (Recent) versus overnight consolidated (Remote) emotional memories, we performed multiple regression analyses on suppression-induced forgetting score separately for Recent and Remote conditions. It is worth noting that suppression-induced forgetting score is quantified by subsequent recall performance for events in the baseline condition contrasting with suppression condition (see **Methods**). The CAR was included as the covariate of interest, while sleep duration, perceived stress, and state-trait anxiety were treated as covariates of no interest. These analyses revealed a significantly positive correlation between CAR and the suppression score in the Recent condition (*r* = 0.336, *p* = 0.002; **Fig. 1C**), but not in the Remote condition (*r* = 0.066, *p* = 0.548; **Fig. 1D**). Further analyses for Fisher’s r-to-z- transformed scores confirmed that the correlation coefficient was significantly higher for the Recent than Remote condition (*Z* = 1.901, *p* = 0.028), indicating an interaction effect in the correlation slopes.

To further characterize the interaction effect between the CAR and memory acquisition time, we conducted a set of complementary analyses by splitting participants into either a Robust- CAR or Lower-CAR group (see **Methods**), based on criteria established in prior studies (Clow et al., 2004; Wust et al., 2000; Xiong et al., 2021). Consistent with above correlational effects, a 2 (Groups: Robust-CAR vs. Lower-CAR) × 2 (acquisition time: Recent vs. Remote) repeated-measures analysis of variance (ANOVA) on suppression-induced forgetting scores revealed confirmed a significant interaction effect between CAR and acquisition time (**Fig. 1G**; *F*_1, 84_ = 5.691, *p* = 0.019, η^2^ = 0.063). Further independent samples t-test confirmed a significant increase in cortisol levels after awakening in the Robust-CAR group compared to the Lower-CAR group (*t*_84_ = 11.383, *p* < 0.001; **Fig. 1E**), but no differences in cortisol levels either before or after fMRI scanning in the afternoon (all *p* > 0.764; **Fig. 1F**). There were no significant group differences in other behavioral and affective measures (all *P* > 0.66; *Fig. S2 & Table S1*). These results indicate that individuals with a Robust-CAR in the morning exhibit greater effectiveness in suppressing recently acquired (i.e., Recent) memories compared to overnight consolidated (i.e., Remote) memories in the afternoon of the same day.

### 3.2 CAR Modulates Neural Activity During Suppression: Prefrontal Engagement and Hippocampal-Striatal Downregulation for Recent Memories

To identify brain systems associated with memory suppression and retrieval for recently acquired versus overnight consolidated memories, we performed a 2 (acquisition time: Recent vs. Remote) × 2 (TNT condition: Think vs. NoThink) repeated-measures ANOVA at the whole-brain level. A significant time × TNT interaction was observed in the bilateral posterior hippocampus and the right amygdala (***Fig. S3***). Specifically, voluntary suppression of recently acquired emotional memories (compared to the Think condition) resulted in weaker activation of these regions (left posterior hippocampus: *F*_1, 66_ = 14.524, *p* < 0.000, η^2^ = 0.180; right posterior hippocampus: *F*_1, 66_ = 8.270, *p* = 0.005, η^2^ = 0.111; right amygdala: *F*_1, 66_ = 4.908, *p* < 0.030, η^2^ = 0.069). Null effects, however, were observed for suppression of overnight emotional memories.

Next, we examined how the CAR proactively modulates functional activation in brain systems involved in suppression-induced forgetting in the afternoon while controlling for sleep duration, perceived stress, and state/trait anxiety. Significant clusters were observed in the bilateral dorsolateral prefrontal cortex (dlPFC) (**Fig. 2A&D**) and middle hippocampus (*Fig. S4A&D; Table S2&S3*). That is, higher CAR was predictive of greater functional activation in these regions during both Recent and Remote conditions (**Fig. 2B&E;** *Fig. S4B&E*). Complementary 2 (acquisition time: Recent vs. Remote) × 2 (group: Robust-CAR vs. Lower-CAR) whole-brain ANOVAs confirmed significant main effects of group in these regions (**Fig. 2C&F**; left dlPFC: *F*_1, 65_ = 4.344, *p* = 0.041, η^2^ = 0.063; right dlPFC: *F*_1, 65_ = 7.282, *p* = 0.009, η^2^ = 0.101).

**Fig. 2.**
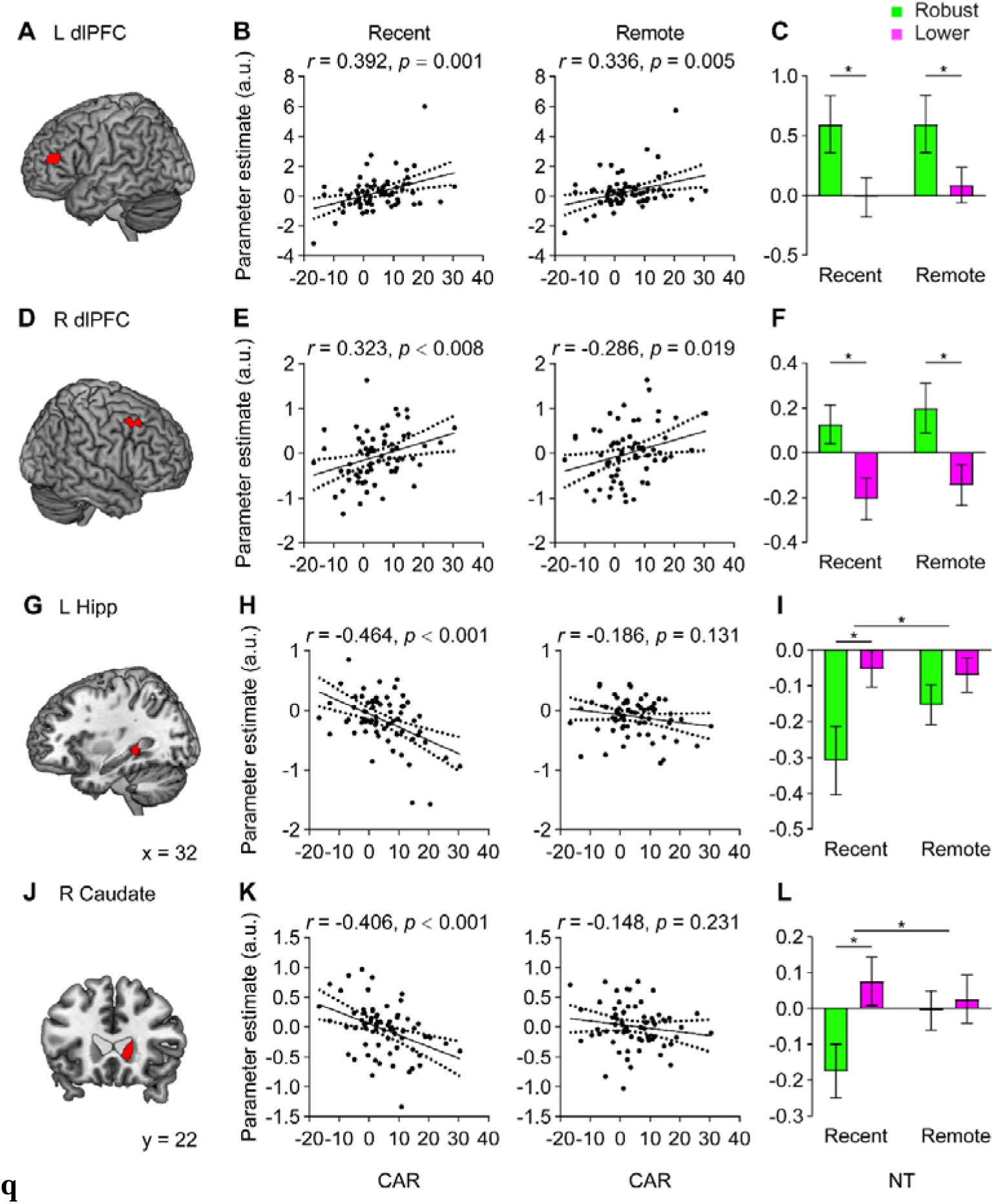
Proactive effects of the CAR on brain systems supporting memory suppression. (**A&D**) Significant clusters in the bilateral dlPFC revealed by whole-brain regression analyses, (**B&E**) with higher CAR predictive of higher functional activation in these regions during both Recent and Remote conditions, (**C&F**) which confirmed by 2-by-2 ANOVAs showing significant main effect of group. (**G&J**) Significant clusters in the left posterior hippocampus and right caudate revealed by whole-brain regression analyses, (**H&K**) with higher CAR predictive of higher functional activation in these regions selectively during Recent but not Remote conditions, (**I&L**) which confirmed by 2-by-2 ANOVAs showing significant group × time interaction effect. Notes are the same as Fig. 2.

Interestingly, significant clusters were also identified in the left posterior hippocampus and right caudate (**Fig. 2G&J**; *Table S2*), wherein higher CAR was predictive of reduced functional activation in these regions during Recent but not Remote condition (**Fig. 2H&K**). Follow-up 2 (acquisition time: Recent vs. Remote) × 2 (group: Robust-CAR vs. Lower-CAR) whole-brain ANOVAs confirmed significant interaction effects in these brain regions (**Fig. 2I&L**; left posterior hippocampus: *F*_1, 65_ = 4.901, *p* = 0.030, η^2^ = 0.070; right caudate: *F*_1, 65_ = 4.557, *p* = 0.037, η^2^ = 0.066). These results indicate that individuals with Robust-CAR exhibit generally greater activation in the prefrontal cortex and middle hippocampus during memory suppression irrespective of acquisition time, but selectively weaker activity in the posterior hippocampus and caudate during suppression of recently acquired emotional memories.

### 3.3 CAR Strengthens Prefrontal-Hippocampal and Prefrontal-Striatal Functional Connectivity During Suppression of Recent Memories

To further investigate how the CAR modulates functional organization of prefrontal, hippocampal, and striatal systems during suppression of recently acquired versus consolidated memories. We performed multiple regression analyses at the whole-brain level for the contrast of suppressing Recent versus Remote memories, with CAR as a covariate of interest, while sleep duration, perceived stress and state-trait anxiety as covariates of no interest.

When using the left posterior hippocampus as a seed region, we identified a significant cluster in the left dlPFC (**Fig. 3A**, *Table S4*), with higher CAR predictive of higher function connectivity during suppression of Recent but not Remote memories (**Fig. 3B**). A follow-up complementary 2 (acquisition time: Recent vs. Remote) × 2 (group: Robust-CAR vs. Lower- CAR) whole-brain ANOVA confirmed a significant interaction effect within this brain system (**Fig. 3C**; *F*_1, 65_ = 8.016, *p* = 0.006, η^2^ = 0.110).

**Fig. 3.**
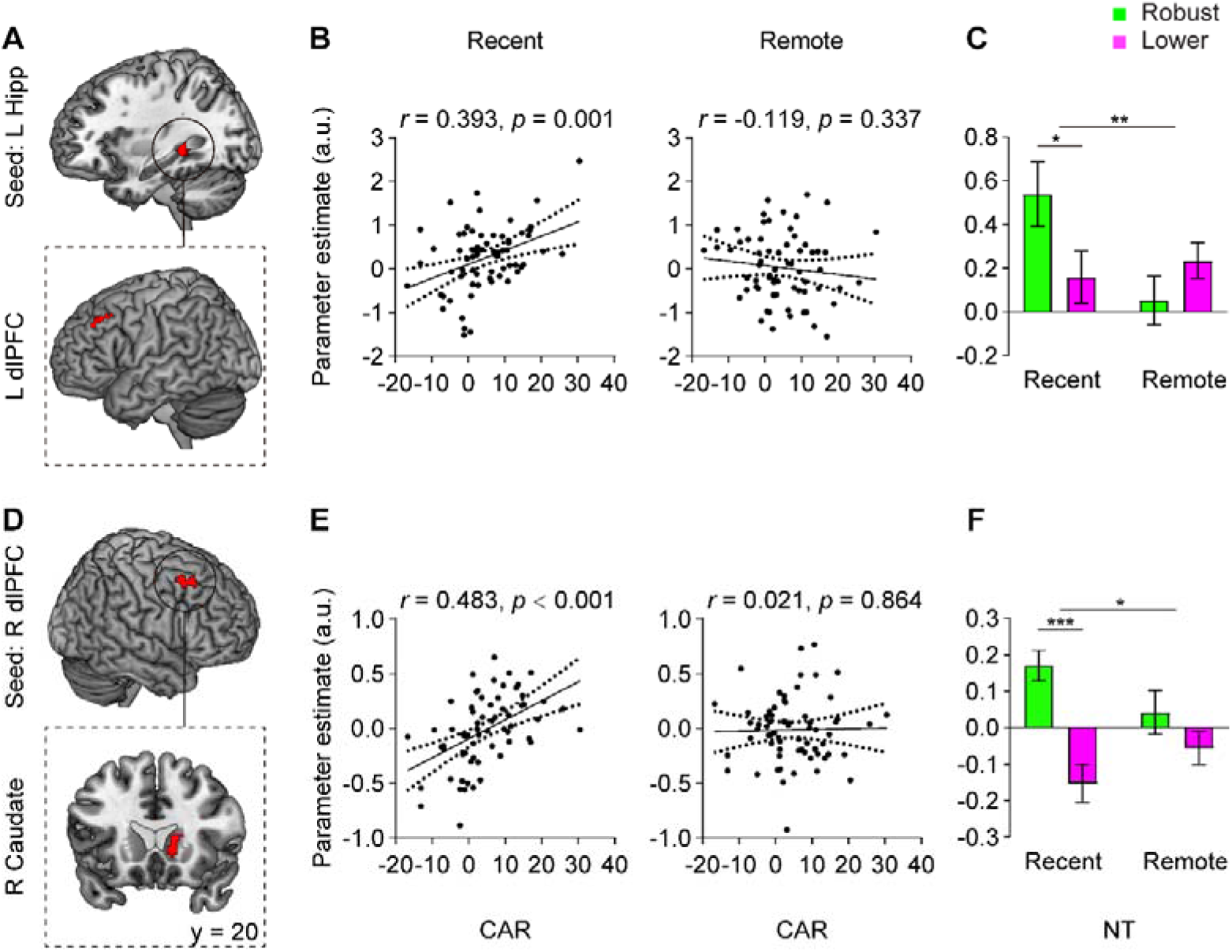
Proactive effects of the CAR on prefrontal coupling with hippocampus and striatal circuitry. (**A&D**) Significant clusters in the left dlPFC revealed by a hippocampal-seeded gPPI analysis and in the right caudate revealed by a dlPFC-seeded gPPI analysis, (**B&E**) with higher CAR predictive of higher functional coupling between these brain systems selectively during Recent but note Remote conditions, (**C&F**) which confirmed by 2-by-2 ANOVAs showing significant group × time interaction effect. Notes are the same as Fig. 2.

When using the right dlPFC as a seed region, we identified significant clusters in the right caudate (**Fig. 3D**, *Table S5*), with higher CAR predictive of greater functional connectivity during suppression of Recent but not Remote memories (**Fig. 3E**). A follow-up 2 (acquisition time: Recent vs. Remote) × 2 (group: Robust-CAR vs. Lower-CAR) whole-brain ANOVA confirmed significant interaction effects within these brain systems (**Fig. 3F**; *F*_1, 65_ = 6.709, *p* = 0.012, η^2^ = 0.094). These results indicate that individuals with robust CAR exhibit greater functional connectivity within prefrontal-hippocampal and prefrontal-caudate circuits selectively during suppression of Recent, but not Remote, emotional memories.

### 3.4 CAR Amplifies Top-Down Effective Connectivity from Prefrontal Cortex to Hippocampal and Striatal Systems During Suppression of Recent Memories

To further investigate how the CAR modulates directional causal influences within prefrontal-hippocampal-striatal circuits during active forgetting of emotional memory, we employed dynamic causal modeling (DCM) supplemented by a hierarchical Parameter Empirical Bayes (PEB) framework (Friston et al., 2003; Zeidman, Jafarian, Corbin, et al., 2019; Zeidman, Jafarian, Seghier, et al., 2019) (**Fig. 4A&B**). This analysis showed that a particular architecture of directed connectivity (Model 8) best explained our data, with a posterior probability of 95%, in relation to alternative plausible architectures (**Fig. 4C**). By thresholding the Bayesian model average (BMA) of directed connectivity estimates at > 95% posterior prob-ability (based on the free energy approximation to the evidence for models with and without connectivity changes), we identified the connections (i.e., model parameters) that were modulated by NoThink_Recent (**Fig. 4D**). These parameters are also shown in **Fig. 4E** as connectivity matrices, in which a positive sign (orange squares) represents a positive modulation while a negative sign (blue squares) represents a negative modulation. Results from BMA showed that there were significant group differences in the modulatory effects of suppression of recently acquired memory on the pathways from the dlPFC to the posterior hippocampus and from the dlPFC to the caudate. Specifically, when actively suppressing emotional memory, individuals with Robust- relative to Lower-CAR group exhibited significantly higher modulatory input for these top-down pathways. These results indicate that individuals with higher CAR exhibit higher top-down effective connectivity from prefrontal to hippocampal-striatal circuit selectively in suppression of Recent but not Remote emotional memories.

**Fig. 4.**
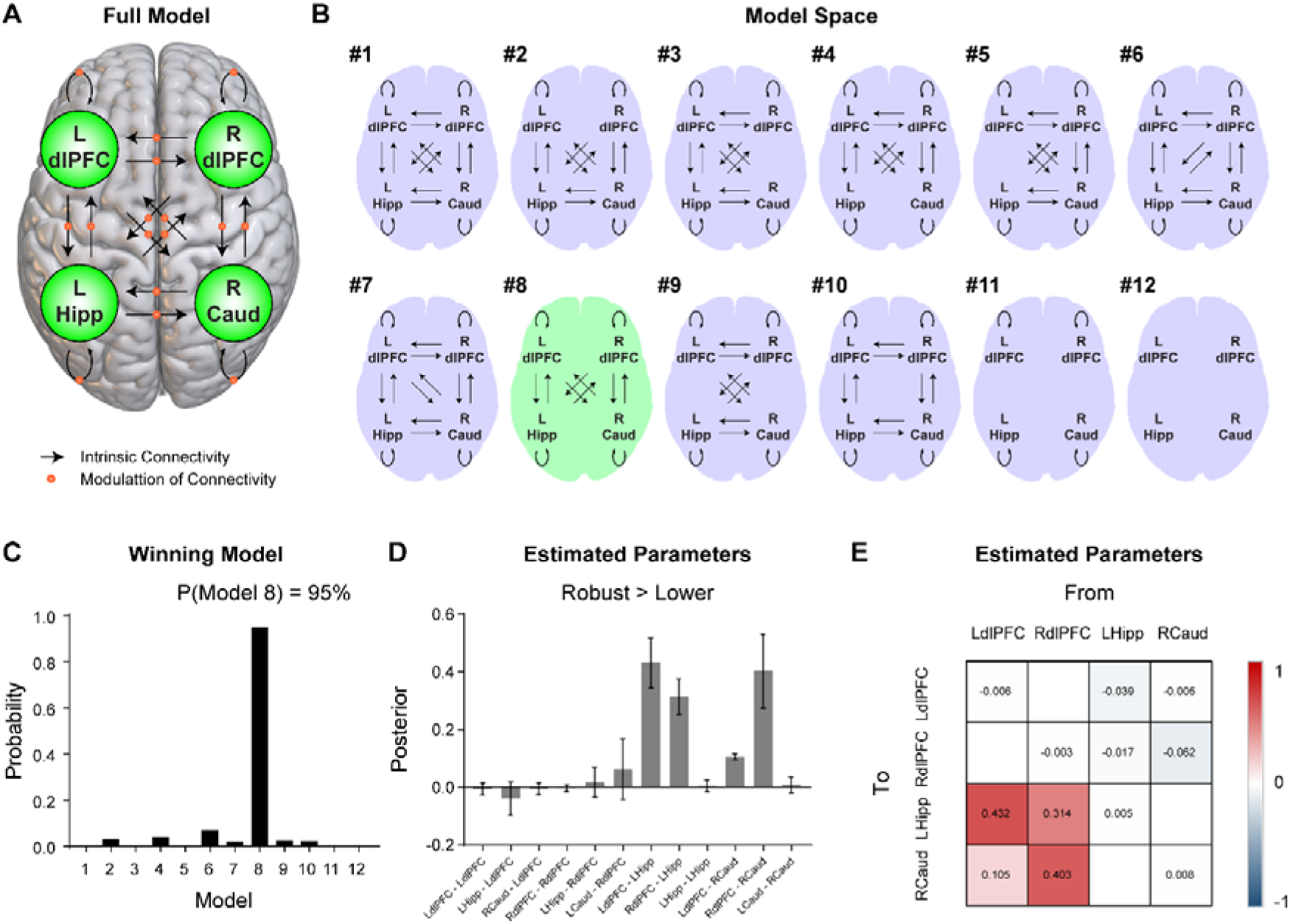
CAR-dependent modulation of top-down effective connectivity from dlPFC to hippocampal-striatal circuits during recent memory suppression. **(A)** Full DCM architecture. Black arrows represent intrinsic connectivity (A-matrix) within the network of four ROIs (bilateral dlPFC, left hippocampus, and right caudate). Red dots represent modulatory effects of NoThink_Recent (i.e., suppression of recently acquired memories) on all connections (B-matrix). **(B)** Model space including the full model, 11 reduced models and a null model. **(C)** The best model was model 8 with the most plausible explanation for the group differences (Robust-CAR vs. Lower-CAR) in terms of changes in effective connectivity modulated by memory suppression. **(D)** BMA of the model parameters survived a thresholding at posterior probability > 95%. Each parameter refers to the modulation effect of NoThink_Recent on each specific connection. **(E)** Effective connectivity matrices.

## 4. Discussion

This study provides compelling evidence for the proactive role of the CAR in modulating the neural circuitry underlying active forgetting, revealing a crucial time-dependent interaction with memory consolidation. Our key findings demonstrate that a robust CAR selectively enhances the efficiency of suppressing recently acquired emotional memories, but not those that have undergone overnight consolidation. This behavioral enhancement is mechanistically underpinned by a dual neurobiological signature: while a robust CAR generally amplifies dlPFC engagement during suppression attempts regardless of memory age, it specifically augments top-down inhibitory control from the dlPFC over the hippocampus and caudate nucleus exclusively for recent traces. These findings significantly advance our understanding of CAR beyond a mere stress biomarker, positioning it as an intrinsic, circadian-entrained orchestrator that proactively synchronizes prefrontal inhibitory networks with the psychophysiological demands of adaptive memory prioritization across sleep-wake cycles.

### 4.1 CAR Proactively and Selectively Facilitates Suppression of Recent Emotional Memories

Behaviorally, our data demonstrates that individuals exhibiting a robust CAR in the morning subsequently displayed significantly greater efficiency in suppressing emotional memories acquired later that same day (Recent), compared to memories consolidated overnight (Remote). This time-dependent effect aligns with the evolutionary perspective embedded within the CAR’s “preparation hypothesis” (Clow et al., 2004; Elder et al., 2014; Fries et al., 2009; Law et al., 2013; Stalder et al., 2025). The CAR is conceptualized as priming the organism for anticipated challenges of the upcoming day, optimizing cognitive and physiological systems for adaptive responses (Chen et al., 2025; Xiong et al., 2021; Zeng et al., 2024). From this vantage point, information encountered during the primed period holds greater immediate relevance and adaptive significance than events from the previous day. Consequently, the cognitive control system, potentially tuned by the CAR’s neuroendocrine milieu, exhibits heightened efficiency in regulating access to these recently encoded, potentially disruptive emotional memories. Our findings extend previous work linking CAR to executive functions like working memory and response inhibition (Shi et al., 2018; Xiong et al., 2021) by demonstrating its specific, proactive influence on the inhibitory control of emotional memories acquired within the same circadian phase. Furthermore, this result resonates with and provides a potential neuroendocrine explanation for our prior observation that consolidated memories become more resistant to suppression(Liu et al., 2016), suggesting that CAR-mediated priming specifically targets memories in their more labile, pre-consolidated state.

### 4.2 CAR Modulates Neural Substrates of Inhibition: Prefrontal Engagement and Hippocampal-Striatal Downregulation

At the neural level, our fMRI results delineate distinct yet complementary mechanisms through which CAR exerts its proactive influence. **First**, we observed that a robust CAR predicted heightened activation within the bilateral dorsolateral prefrontal cortex (dlPFC) during suppression attempts for both Recent and Remote memories. This finding underscores a generalized role for CAR in potentiating the engagement of this key cognitive control hub, consistent with the dlPFC’s established function in implementing inhibitory control over memory retrieval (Anderson et al., 2004; Benoit & Anderson, 2012). This broad amplification of dlPFC activity suggests that CAR establishes a baseline “preparatory tone” within prefrontal executive systems, enhancing their readiness to exert control irrespective of the specific temporal characteristics of the targeted memory.

**Second**, and more critically, CAR exhibited a time-specific modulatory effect on downstream limbic and striatal regions. Individuals with a robust CAR showed significantly reduced activation specifically in the left posterior hippocampus and right caudate during suppression of Recent memories. This pattern of regionally specific down regulation is consistent with the neural signature of successful memory suppression: dlPFC-mediated inhibition leading to the dampening of hippocampal retrieval processes (Anderson & Hanslmayr, 2014; Depue et al., 2016) and striatal involvement in supramodal inhibitory control (Burke et al., 2017; Guo et al., 2018; Robbins & Cools, 2014). The dissociation between middle (central) hippocampal regions showing general CAR-related modulation and posterior hippocampal regions showing time-specific suppression-related deactivation is particularly noteworthy. This aligns with proposed functional specializations within the hippocampus, where posterior regions may be more critically involved in detailed, context-specific memory representations and retrieval (Poppenk & Moscovitch, 2011), making them a primary target for inhibitory control when suppressing recent, potentially vivid traces. The caudate nucleus, implicated as a hub for translating prefrontal control signals into action suppression (Guo et al., 2018; R. Levy et al., 1997), also demonstrates CAR-dependent downregulation specifically for recent memories, highlighting the striatum’s crucial role in this inhibitory circuit. Thus, CAR’s selective effect on suppression efficiency for recent memories appears to be mechanistically implemented through its specific amplification of inhibitory pathways targeting the posterior hippocampus and caudate.

### 4.3 CAR Enhances Functional and Effective Connectivity Within Inhibitory Circuits for Recent Memories

#### Functional Connectivity Signatures

Moving beyond regional activation, our connectivity analyses provide further mechanistic insight into how CAR facilitates inhibitory control through network-level reorganization. This approach aligns with established frameworks for investigating memory control circuits (Anderson & Hanslmayr, 2014; Depue et al., 2016). Using the left posterior hippocampus as a seed region, we observed that stronger CAR predicted increased functional coupling between this region and the left dlPFC specifically during suppression of Recent memories. This finding resonates with prior evidence that prefrontal-hippocampal functional connectivity mediates memory suppression success (Benoit, Hulbert, Huddleston, & Anderson, 2015; Gagnepain, Hulbert, & Anderson, 2017). Similarly, when seeding the right dlPFC, stronger CAR predicted enhanced functional coupling with the right caudate nucleus—a key striatal region implicated in translating control signals into action suppression (Guo et al., 2018; R. Levy et al., 1997)—again selectively for Recent suppression. This pattern indicates that CAR not only modulates activity levels within key nodes but also strengthens their functional integration specifically when suppressing recently acquired emotional content.

#### Directed Effective Connectivity

Crucially, dynamic causal modeling (DCM) with Parametric Empirical Bayes (Zeidman, Jafarian, Corbin, et al., 2019; Zeidman, Jafarian, Seghier, et al., 2019) provided direct evidence for CAR’s influence on directional information flow. Our model comparison identified a specific architecture where group differences (Robust- vs. Lower-CAR) were best explained by modulation of directed connectivity pathways. Bayesian Model Averaging revealed significantly stronger top-down modulatory influences from dlPFC to both posterior hippocampus and caudate nucleus during recent memory suppression in Robust-CAR individuals. This finding is pivotal, as it demonstrates that CAR proactively enhances the effective connectivity underlying the dlPFC’s inhibitory control over two critical systems: the hippocampal retrieval engine (Anderson & Hulbert, 2021) and the striatal action-suppression hub (Robbins & Cools, 2014). It directly supports the hypothesis that CAR’s preparatory role involves strengthening the very pathways responsible for implementing memory suppression, particularly for memories acquired within the same circadian window. This enhanced top-down control likely facilitates the rapid and efficient gating or inhibition of hippocampal retrieval processes and striatal responses, leading to the observed behavioral advantage in suppressing recent emotional memories.

### 4.4 Toward a Model of CAR-Mediated Neuroendocrine Pre-tuning for Adaptive Forgetting

Integrating our behavioral, univariate fMRI, functional connectivity, and effective connectivity results, we propose a model of “CAR-mediated neuroendocrine pre-tuning” for adaptive memory control. In this model, the CAR surge upon awakening serves as an endogenous, circadian-timed signal that proactively establishes a physiological and neural state optimized for cognitive control demands anticipated during the day. This pre-tuning manifests at the neural level through two synergistic mechanisms: (1) a generalized potentiation of dlPFC readiness for cognitive control operations, and (2) a specific sensitization or strengthening of the top-down inhibitory pathways (dlPFC to Hippocampus and dlPFC to Caudate) responsible for suppressing unwanted, recently encoded emotional information. This pre-tuning aligns with glucocorticoid receptor (GR) mediated genomic actions in the brain, which can prime neural circuits over slower time courses (de Kloet et al., 2019; Herman et al., 2003). The CAR might thus initiate genomic cascades that enhance the excitability and connectivity of prefrontal inhibitory networks and their projections to limbic/striatal targets, setting the stage for more effective phasic inhibitory control when needed later in the day. This model provides a neuroendocrine explanation for the time- sensitive window of enhanced suppression efficacy we observed and bridges circadian biology with cognitive neuroscience models of memory control.

### 4.5 Implications and Limitations

Our findings have significant implications for understanding and treating stress-related psychiatric disorders characterized by intrusive memories and circadian dysregulation (e.g., PTSD, depression). The identification of CAR as a key modulator of active forgetting circuitry suggests that strategies aimed at stabilizing or enhancing CAR (e.g., chronotherapy, light therapy, behavioral routines) might offer novel avenues for therapeutic intervention, particularly for disorders involving maladaptive recollection of recent traumatic events. Furthermore, understanding the time-dependent vulnerability of memories to suppression, modulated by CAR, could inform the timing of therapeutic interventions like exposure therapy or cognitive control training.

Several limitations warrant consideration. First, our sample consisted exclusively of young, healthy males to control for sex hormone influences. While this enhances internal validity, it limits the generalizability of our findings. Future research must investigate whether these mechanisms operate similarly in females and across different age groups or clinical populations.

Second, the correlational nature of our design precludes definitive causal claims about CAR’s influence. Pharmacological manipulation of cortisol levels or CAR, or naturalistic studies monitoring CAR fluctuations and suppression ability longitudinally, are needed to establish causality. Third, while we controlled for several potential confounds (sleep duration, anxiety, stress), other factors like stress anticipation or sleep quality details could influence results and merit closer examination. Fourth, the functional dissociation between hippocampal subregions (middle vs. posterior) requires further validation using higher-resolution imaging techniques.

### 4.6 Conclusion

In conclusion, our study provides robust evidence that the CAR plays a proactive and time- specific role in facilitating the active forgetting of unwanted emotional memories. By demonstrating that a robust CAR selectively enhances suppression efficiency for recently acquired memories through a dual mechanism of heightened dlPFC engagement and amplified top-down inhibitory control over hippocampal-striatal circuitry, we refine the functional significance of CAR from a passive stress indicator to an active, circadian- entrained orchestrator of adaptive memory prioritization. These findings bridge fundamental mechanisms of circadian biology, cognitive neuroscience, and clinical psychopathology, paving the way for novel neuroendocrine and chronotherapeutic strategies aimed at alleviating maladaptive recollection in stress-related mental disorders.

## Supporting information

Supplemental Informations

## Acknowledgements

This study has been supported by the Scientific and Technological Innovation 2030-Major Projects 2021ZD0200500, the National Natural Science Foundation of China (32130045, 32361163611, and 82021004), the Major Project of National Social Science Foundation (19ZDA363 and 20&ZD153), and the Fundamental Research Funds for the Central Universities (310422131 and 310400209508).

