## Supplemental Informations for "Brain Preparedness for Active Forgetting: Cortisol Awakening Response Proacts Prefrontal Control Over Hippocampal-Striatal Circuitry"

Supplementary Figures

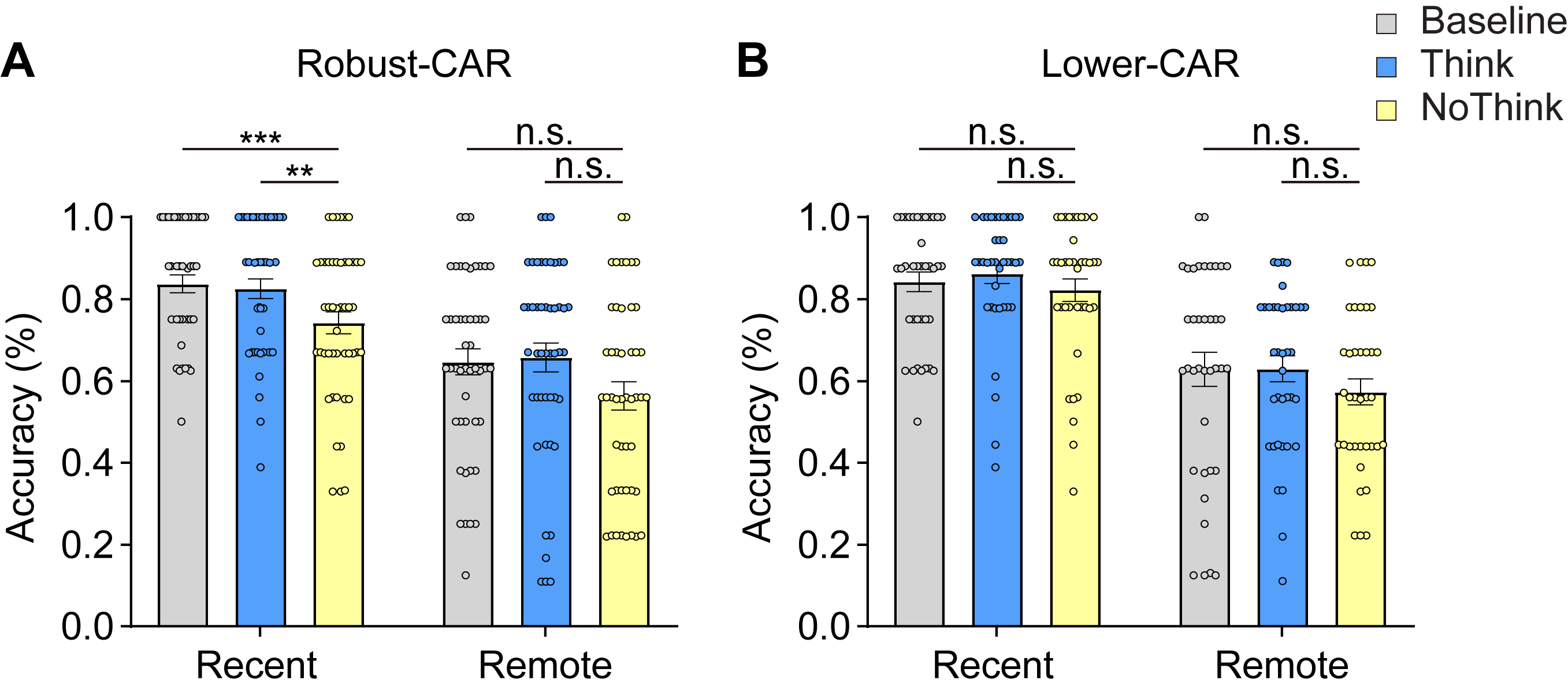

### **Fig. S1. Behavioural performance.**

(**A-B**) Bar graphs depict cued-recall accuracy for Recent and Remote memories as a function of ‘Baseline’, ‘Think’ and ‘NoThink’ during the test phase for Robust- and Lower-CAR group, respectively. Notes: n.s., not significant; *, *p* < 0.05; **, *p* < 0.01; ***, *p* < 0.001; Error bars represent standard error of mean.

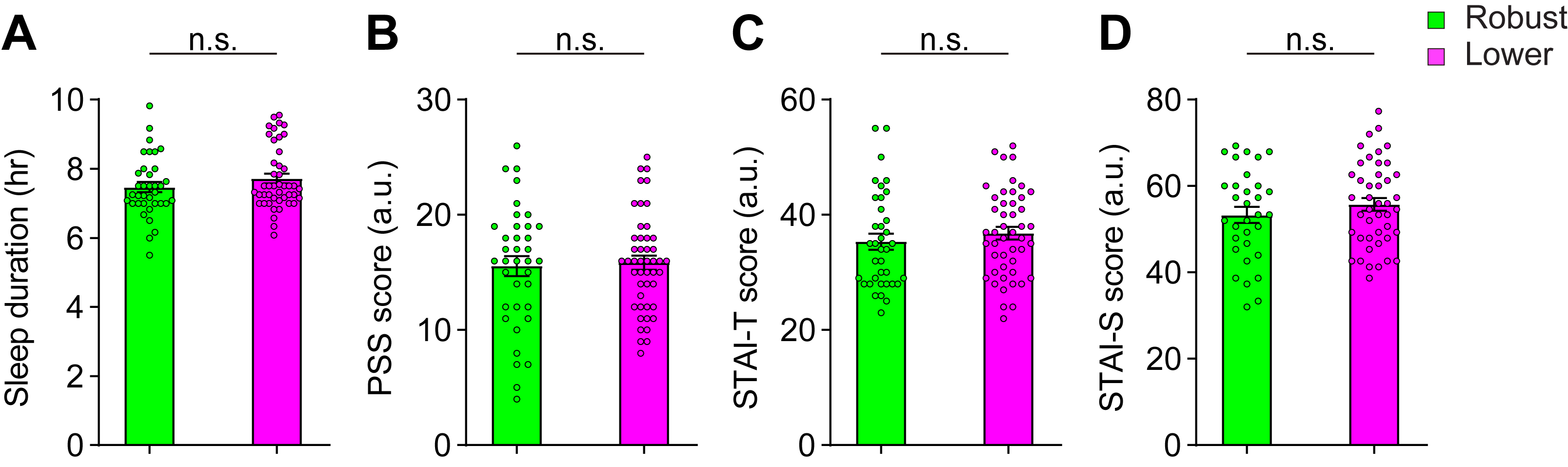

### **Fig. S2. Psychological measurement.**

Bar graphs depict that there was no significant group difference for (**A**) sleep duration, (**B**) PSS, (**C**) STAI-S and (**D**) STAI-T scores (**Table S2**). Notes are the same as Fig. S1

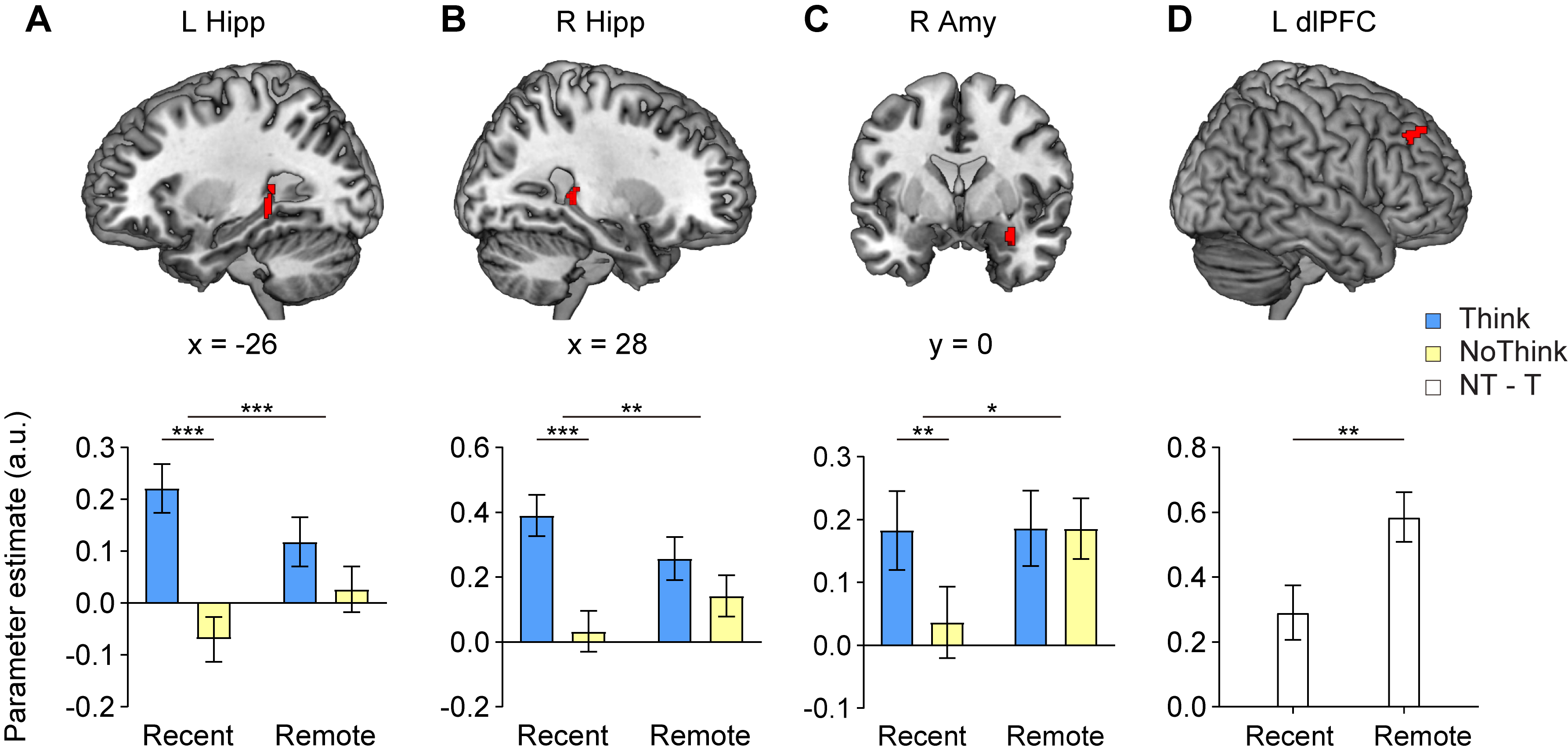

### **Fig. S3. Brain systems involved in active suppression of emotional memory.** Brain activation in the bilateral posterior hippocampus (Hipp; **A-B**) and the right amygdala (Amy; **C**) was lower in active suppression (Nothink) relative to recall (Think) newly acquired (Recent) emotional memories, while no such difference was found in overnight-consolidated (Remote) memories. Notes are the same as Fig. S1.

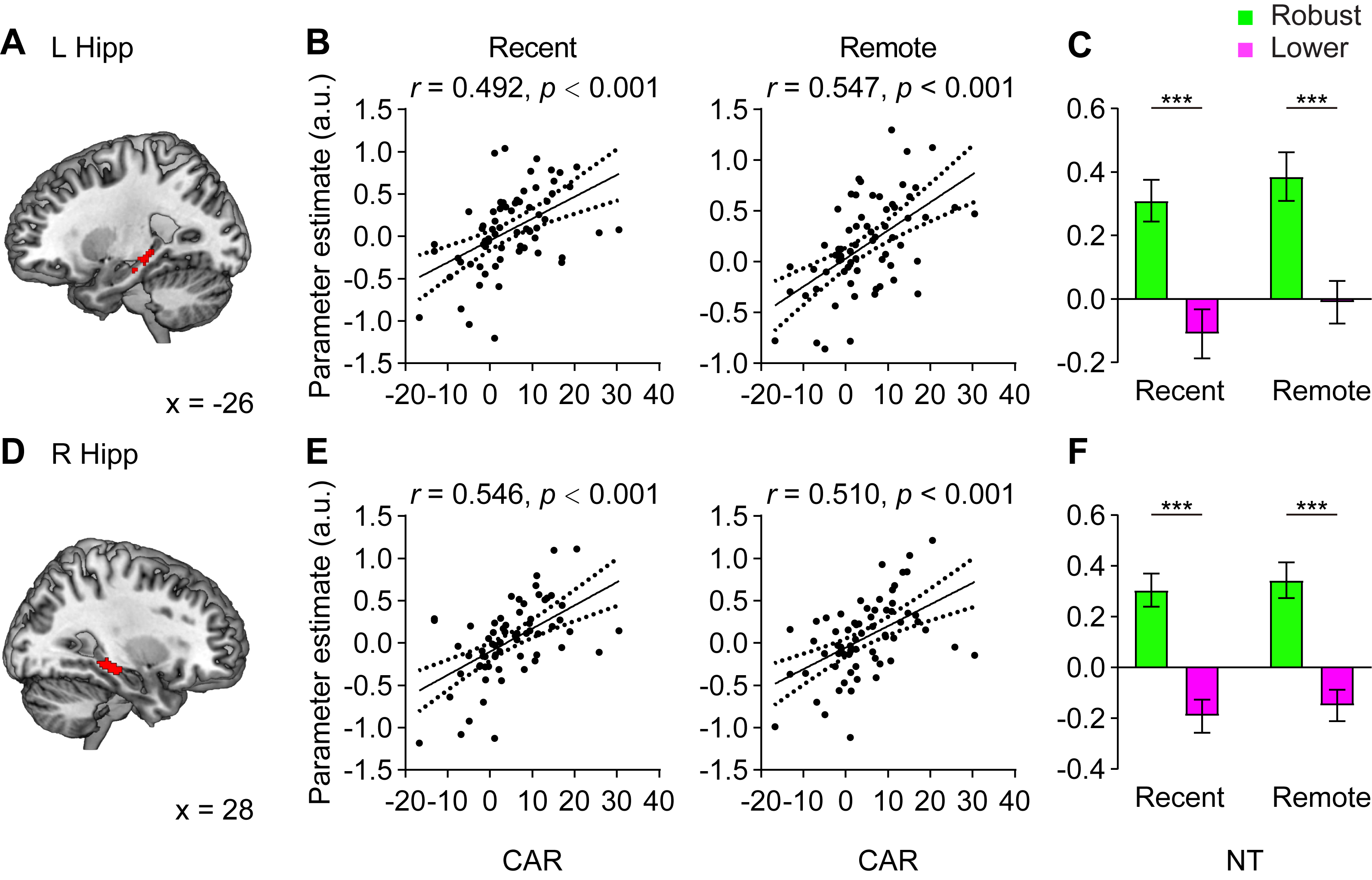

### **Fig. S4. Proactive effects of the CAR on brain systems supporting memory suppression.** (**A&D**) Significant clusters in the bilateral middle hippocampus revealed by whole-brain regression analyses, (**B&E**) with higher CAR predictive of higher functional activation in these regions during both Recent and Remote conditions, (**C&F**) which confirmed by 2-by-2 ANOVAs showing significant main effect of group. Notes are the same as Fig. S1.

### **Supplementary Tables**

### **Table S1. Participant** **demographics and psychological measurements**

|  | **Behavior** | | | **Brain** | | |
| --- | --- | --- | --- | --- | --- | --- |
|  | Robust CAR | Lower CAR | *p* | Robust CAR | Lower CAR | *p* |
| N | 46 | 40 |  | 29 | 38 |  |
| Age | 22.19±1.20 | 21.96±1.43 | 0.43 | 22.17±1.20 | 22.03±1.37 | 0.65 |
| (Range) | (21-25) | (18-26) |  | (21-25) | (18-26) |  |
| Sleep duration | 7.47±0.88 | 7.65±1.02 | 0.40 | 7.38±0.74 | 7.66±1.10 | 0.26 |
| PSS | 15.57±5.31 | 15.85±4.22 | 0.79 | 15.76±5.48 | 16.03±4.23 | 0.82 |
| STAI_S | 35.35±8.37 | 36.81±7.53 | 0.41 | 36.48±8.73 | 36.79±7.09 | 0.87 |
| STAI_T | 39.97±7.95 | 41.84±7.37 | 0.29 | 40.24±7.84 | 41.97±7.67 | 0.39 |

Mean (± standard deviation) of demographics and psychological measurements. P values represent the significance of comparisons between groups. Notes: N, participants numbers; PSS, perceived stress scale; STAI-S, state trait anxiety inventory-state; STAI-T, state trait anxiety inventory-trait.

### **Table S2. Brain regions modulated by the CAR in suppression of recent memory**

| **Region** | **L/R** | **BA** | **MNI (x, y, z)** | | | **T** |
| --- | --- | --- | --- | --- | --- | --- |
| **Regression (Positive effects): CAR - (Recent NT vs. fixation)** | | | | | | |
| **Hippocampus** | R | 28 | 34 | -26 | -12 | 5.40 |
|  |  |  | 34 | -16 | -6 | 4.22 |
|  |  |  | 30 | -8 | -10 | 3.73 |
| Thalamus | L | - | -22 | -20 | 0 | 4.91 |
|  |  |  | -26 | -30 | 0 | 3.62 |
| Supra Marginal | L | 2, 4, 3, 40 | -54 | -20 | 32 | 4.72 |
| Cingulum Mid | L | 24 | -8 | -2 | 34 | 4.48 |
| **Frontal Mid** | R | 8, 6 | 28 | 12 | 50 | 4.37 |
| Insula | L | 13, 22 | -36 | -18 | 4 | 4.21 |
|  |  |  | -32 | -16 | -4 | 3.87 |
|  |  |  | -30 | -6 | -6 | 3.75 |
| Calcarine | R | 30, 17, 18, 23 | 16 | -70 | 8 | 4.15 |
| Insula | L | 13 | -34 | -4 | 14 | 3.95 |
|  |  |  | -24 | -6 | 16 | 3.41 |
| **Regression (Negative effects): CAR - (Recent NT vs. fixation)** | | | | | | |
| **Caudate** | L | - | -8 | 24 | 2 | 5.01 |
|  |  |  | -14 | 24 | 8 | 4.99 |
|  |  |  | -16 | 30 | 0 | 3.77 |
| Caudate | R | - | 16 | 28 | 12 | 4.35 |
|  |  |  | 14 | 28 | 2 | 4.15 |

Only clusters, significant at a height threshold of *P* < 0.001 and an extent threshold of *P* < 0.05 corrected on the whole brain level, are reported with local maxima in Montreal Neurological Institute (MNI) space. Clusters in the **Hippocampus, Middle Frontal Gyrus and Caudate** regions are in bold. Notes: NT, no think; L, left hemisphere; R, right hemisphere; BA, Brodmann’s area.

### **Table S3. Brain regions modulated by the CAR in suppression of remote memory**

| **Region** | **LR** | **BA** | **MNI (x, y, z)** | | | **T** |
| --- | --- | --- | --- | --- | --- | --- |
| **Regression (Positive effects): CAR - (Remote NT vs. fixation)** | | | | | | |
| **Hippocampus** | R | 28, 35 | 34 | -26 | -12 | 5.70 |
|  |  |  | 34 | -16 | -4 | 4.75 |
|  |  |  | 32 | -20 | 4 | 4.52 |
| Insula | L | 13, 21, 28, 35, 22 | -34 | -18 | -2 | 5.24 |
|  |  |  | -18 | -24 | -6 | 4.99 |
|  |  |  | -24 | -20 | 0 | 4.45 |
| Cingulum_Mid | L | 31 | -16 | -28 | 40 | 4.78 |
| Postcentral | L | 2, 3, 40, 4 | -52 | -18 | 30 | 4.58 |
|  |  |  | -56 | -22 | 36 | 3.83 |
| Calcarine | R | 30, 18, 17, 23, 31, 19 | 18 | -76 | 8 | 4.46 |
| Cingulum_Mid | L | 24, 32 | -8 | 0 | 38 | 4.11 |
|  |  |  | -2 | -2 | 44 | 3.31 |
| **Frontal_Mid** | R | 8 | 28 | 14 | 46 | 4.07 |

### **Table S4. Hippocampus-seeded functional coupling between modulated by the CAR in suppression of remote memory**

| **Region** | **LR** | **BA** | **MNI** | | | **T** |
| --- | --- | --- | --- | --- | --- | --- |
| **Regression (Positive effects): CAR - (Recent NT vs. fixation)** | | | | | | |
| Insula | R | - | 44 | 2 | 6 | 4.24 |
| Cingulum_Mid | R | 31 | 18 | -32 | 46 | 3.72 |
|  |  |  | 26 | -36 | 44 | 2.71 |
| **Putamen** | R |  | 30 | -8 | 6 | 3.72 |
| ParaHippocampal | R |  | 22 | -42 | -2 | 3.60 |
|  |  |  | 24 | -30 | -2 | 3.16 |
|  |  |  | 30 | -38 | -6 | 3.11 |
| Angular | R | 39 | 52 | -60 | 24 | 3.54 |
| Insula | LR | 13, 44, 22 | -46 | 0 | 4 | 3.43 |
|  |  |  | -40 | 12 | 4 | 2.98 |
| Frontal_Sup | L | 6, 4 | -20 | -8 | 66 | 3.13 |
|  |  |  | -20 | -24 | 60 | 3.08 |
|  |  |  | -18 | -16 | 66 | 2.99 |

### **Table S5. MFG-seeded functional coupling between modulated by the CAR in suppression of remote memory**

| **Region** | **LR** | **BA** | **MNI** | | | **T** |
| --- | --- | --- | --- | --- | --- | --- |
| **Regression (Positive effects): CAR - (Recent NT vs. fixation)** | | | | | | |
| Insula | R | - | 44 | 2 | 6 | 4.24 |
| Cingulum_Mid | R | 31 | 18 | -32 | 46 | 3.72 |
|  |  |  | 26 | -36 | 44 | 2.71 |
| Putamen | R |  | 30 | -8 | 6 | 3.72 |
| ParaHippocampal | R |  | 22 | -42 | -2 | 3.60 |
|  |  |  | 24 | -30 | -2 | 3.16 |
|  |  |  | 30 | -38 | -6 | 3.11 |
| Angular | R | 39 | 52 | -60 | 24 | 3.54 |
| Insula | LR | 13, 44, 22 | -46 | 0 | 4 | 3.43 |
|  |  |  | -40 | 12 | 4 | 2.98 |
| Frontal_Sup | L | 6, 4 | -20 | -8 | 66 | 3.13 |
|  |  |  | -20 | -24 | 60 | 3.08 |
|  |  |  | -18 | -16 | 66 | 2.99 |
